# Intermittent hypoxia enhances the expression of HIF1A by increasing the quantity and catalytic activity of KDM4A-C and demethylating H3K9me3 at the *HIF1A* locus

**DOI:** 10.1101/2021.07.25.453726

**Authors:** Chloe-Anne Martinez, Neha Bal, Peter A Cistulli, Kristina M Cook

## Abstract

Cellular oxygen-sensing pathways are primarily regulated by hypoxia inducible factor-1 (HIF-1) in chronic hypoxia and are well studied. Intermittent hypoxia also occurs in many pathological conditions, yet little is known about its biological effects. In this study, we investigated how two proposed cellular oxygen sensing systems, HIF-1 and KDM4A-C, respond to cells exposed to intermittent hypoxia and compared to chronic hypoxia. We found that intermittent hypoxia increases HIF-1 activity through a pathway distinct from chronic hypoxia, involving the KDM4A, -B and -C histone lysine demethylases. Intermittent hypoxia increases the quantity and activity of KDM4A-C resulting in a decrease in H3K9 methylation. This contrasts with chronic hypoxia, which decreases KDM4A-C activity, leading to hypermethylation of H3K9. Demethylation of histones bound to the *HIF1A* gene in intermittent hypoxia increases *HIF1A* mRNA expression, which has the downstream effect of increasing overall HIF-1 activity and expression of HIF target genes. This study highlights how multiple oxygen-sensing pathways can interact to regulate and fine tune the cellular hypoxic response depending on the period and length of hypoxia.

## Introduction

Chronic hypoxia is common in solid tumors and is associated with treatment resistance and aggressive disease (Semenza, 2012). Along with chronically hypoxic regions, solid tumors contain regions of intermittent hypoxia, which arise from fluctuations in perfusion and red blood cell flux (Saxena & Jolly, 2019). Systemic intermittent hypoxia also occurs in obstructive sleep apnea (OSA), a common disorder characterized by repetitive interruptions in breathing during sleep due to recurrent collapse of the upper airway. Like chronic hypoxia, perfusion-limited intermittent hypoxia and OSA-induced intermittent hypoxia have been shown to promote a more malignant tumor phenotype (Cardenas-Navia et al., 2008; Hunyor & Cook, 2018). However, when compared to chronic hypoxia, the molecular mechanisms driving cell behavior in intermittent hypoxia are less well understood.

The cellular response to chronic hypoxia is largely coordinated by hypoxia inducible factors (HIFs). HIF is controlled by the prolyl hydroxylase (PHD) oxygen-sensing enzymes, which use oxygen to hydroxylate HIF-α subunits, targeting them for degradation. In the presence of insufficient oxygen, PHD activity substantially decreases, enabling HIF-α to bind to HIF-β and initiate the transcription of target genes involved in metabolism, angiogenesis, cell survival, and apoptosis. Similar to chronic hypoxia, longer periods of intermittent hypoxia (hours to days) have been shown to increase HIF-1 activity (Hunyor & Cook, 2018).

Much less is known about the HIF response to rapid intermittent hypoxia (minutes) as occurs in sleep apnea, though there have been a series of studies indicating HIF1-α increases in the carotid body in intermittent hypoxia. HIF-1α increases were through Ca^2+^- and protein kinase C-dependent activation of mTOR, which increases HIF-1α protein synthesis. Furthermore, HIF-1α degradation decreased from impaired prolyl hydroxylation. In contrast, HIF-2α decreased in the carotid body in intermittent hypoxia through increased degradation by Ca2+-activated calpain proteases (Prabhakar & Semenza, 2016).

Epigenetic modifications are also involved in the response to chronic hypoxia (Hancock, Dunne, Walport, Flashman, & Kawamura, 2015). Histone lysine demethylases, known as KDMs (or JmjCs), remove methyl groups from histone lysine residues. The methylation status of histones affects the chromatin structure, and alters accessibility of chromatin to transcription factors and therefore the expression of genes (Kouzarides, 2007). Similar to the PHDs, which regulate HIF-α degradation, KDM activity requires 2-oxoglutarate, Fe(II) and oxygen as essential cofactors (Hancock et al., 2015). Specific members of the KDM family have been found to act as enzymatic oxygen sensors including KDM4A (Hancock, Masson, Dunne, Flashman, & Kawamura, 2017), KDM5A (Batie et al., 2019) and KDM6A (Chakraborty et al., 2019), meaning that their activity is affected by changes in physiologically relevant oxygen levels. Chronic hypoxia inhibits KDM activity resulting in global hypermethylation of histone lysine residues, altering the expression of several genes (Hancock et al., 2015). To our knowledge, no one has examined how KDMs behave in intermittent hypoxia, nor their interaction with the HIF pathway under these conditions.

In this study, we describe a unique role for the KDM4A, KDM4B and KDM4C histone demethylases in regulating the expression of *HIF1A*, ultimately controlling the HIF-1 response in rapid intermittent hypoxia. This pathway is distinct from the canonical oxygen-dependent degradation pathway and *does not occur in chronic hypoxia*.

## Results

### Intermittent hypoxia increases HIF-1α protein and expression of known HIF-1 target genes

We have previously shown that HIF-1α protein and HIF-1 target gene expression increases in intermittent hypoxia in HCT116 cells (C. Martinez, Kerr, Jin, Cistulli, & Cook, 2019). To examine if this is a generalized cellular response, we exposed MCF7, MDA-MB-231, and PC3 cells to rapid intermittent hypoxia and compared the results to normoxic and chronically hypoxic conditions. Relative to normoxia, HIF-1α protein increased under both chronic and intermittent hypoxia, but to a lesser extent in intermittent hypoxia, similar to HCT116 cells (C. Martinez et al., 2019) (Fig 1A). We also measured the expression of HIF-1 target genes involved in glycolysis, extracellular matrix (ECM) remodeling and the HIF pathway. Expression for *SLC2A1* (GLUT1, glucose transporter type-1), *LDHA* (lactate dehydrogenase A), *PLOD2* (procollagen-lysine,2-oxoglutarate 5- dioxygenase 2), *P4HA1* (prolyl 4-hydroxylase subunit alpha 1), *P4HA2* (prolyl 4-hydroxylase subunit alpha 2), *EGLN1* (PHD2, prolyl hydroxylase domain 2), and *EGLN3* (PHD3, prolyl hydroxylase domain 3) increased in all tested cell lines exposed to chronic or intermittent hypoxia (Fig 1B-D). Interestingly, *HK2* (hexokinase 2) decreased in MDA-MB-231 cells exposed to intermittent hypoxia (Fig 1C), while increasing in MCF7 and PC3 cells (Fig 1B and D). The reasons for these discrepancies are not clear but may be due to cell line-specific gene regulation differences or mutations. Overall, the increase in HIF-1α and HIF target gene expression in chronic and intermittent hypoxia supports the idea that HIF-1 is activated as a general response to both forms of hypoxia, and as we showed previously, the degree of gene expression is related to a hypoxia dose- dependent effect (C. Martinez et al., 2019).

**Figure 1:**
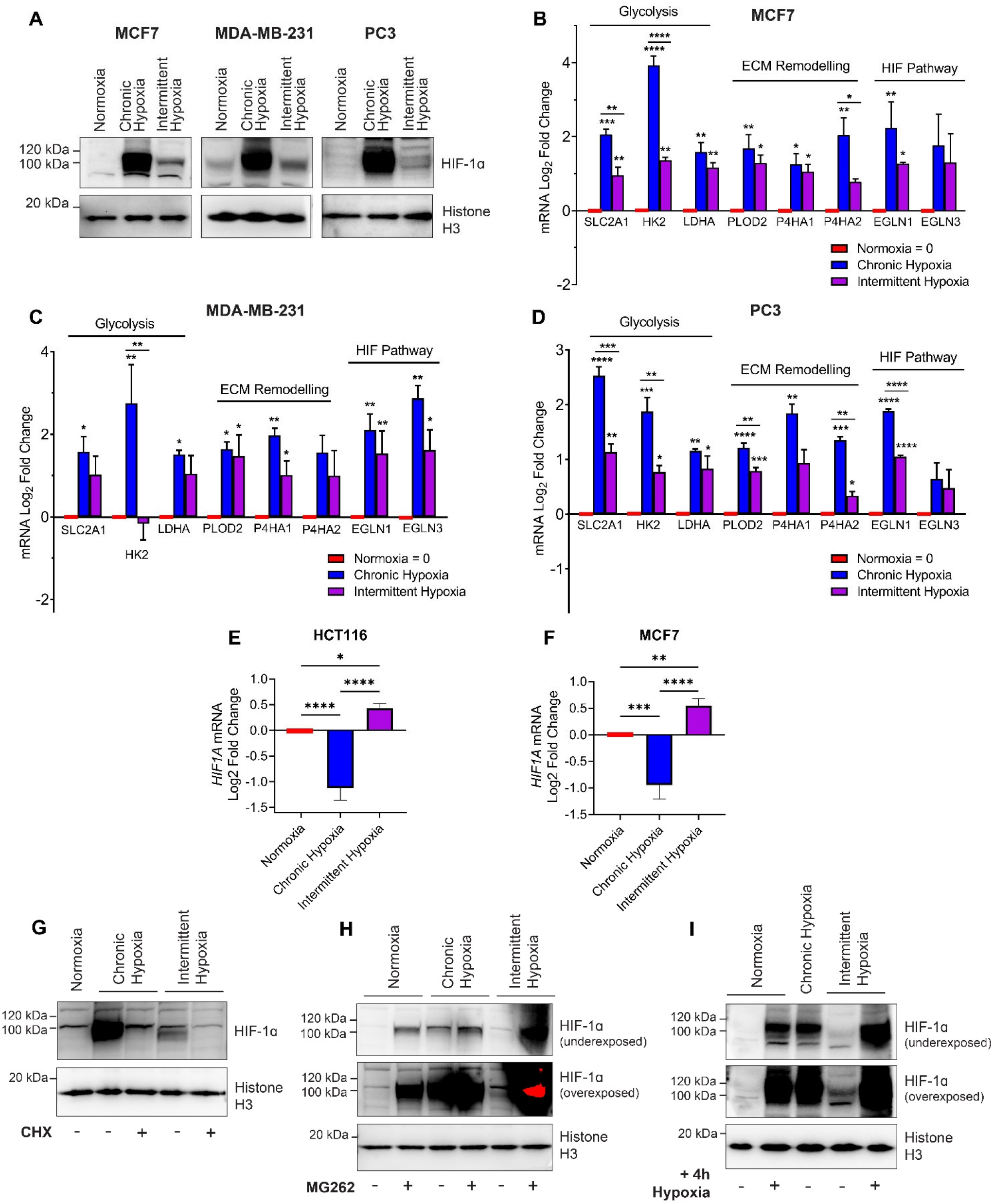
Intermittent hypoxia activates HIF-1 through increased expression of *HIF1A*. (A) Nuclear HIF-1α in MCF7, MDA- MB-231, and PC3 cells following exposure to normoxia, chronic hypoxia and intermittent hypoxia over 18 h. (B-D) mRNA expression of HIF-1 target genes involved in glycolysis (*SLC2A1, HK2, LDHA*), extracellular matrix remodeling (*PLOD2, P4HA1, P4HA2*) and the HIF pathway (*EGLN1, EGLN3*) in MCF7 (B), MDA-MB-231 (C), and PC3 (D) cells after 18 h exposure to oxygen conditions. All values are normalized to normoxic expression levels which is set to 0 on the Log2 scale. Results are the mean ± S.E.M of 3 independent experiments run in duplicate. (E) *HIF1A* mRNA levels in HCT116 cells and MCF7 cells (F) after 18 h exposure to normoxia, chronic hypoxia and intermittent hypoxia. Results are the mean ± S.E.M of 3 independent experiments. ^*^ p < 0.05, ^**^ p < 0.01, ^***^ p < 0.001, ^****^ p < 0.0001. Asterisks above a data point (without line) indicate significance as compared to normoxia. Asterisks above a line compare data points connected by the line. (G) Nuclear HIF-1α protein levels in HCT116 cells exposed to normoxia, chronic hypoxia and intermittent hypoxia treated with DMSO or 10 μg/ml cycloheximide (CHX, protein synthesis inhibitor). (H) Nuclear HIF-1α in HCT116 cells exposed to normoxia, chronic hypoxia and intermittent hypoxia with DMSO or 1 µM MG262 (proteasome inhibitor). (I) Nuclear HIF- 1α in MCF7 cells exposed to 18 h normoxia, chronic hypoxia, and intermittent hypoxia with or without an additional 4 h exposure to chronic hypoxia. Nuclear histone H3 is used as a loading control for (A), (G), (H) and (I).

### Intermittent hypoxia increases *HIF1A* mRNA

HIF-1α protein is degraded in normoxia through an oxygen- dependent mechanism and in hypoxia, HIF-1α protein is stabilized. Expression of the *HIF1A* gene is believed to be constitutive (Wenger, Rolfs, Marti, Guénet, & Gassmann, 1996), and many studies have focused on the post-translational regulation of HIF-1α with considerably less attention on expression of the *HIF1A* gene under different oxygen conditions. Having identified that HIF-1 activity increases in a wide range of cells exposed to intermittent hypoxia, we wanted to know if *HIF1A* mRNA expression was affected by intermittent hypoxia or if regulation of HIF-1α protein levels was occurring primarily through the oxygen-dependent post- translational modifications pathway as in chronic hypoxia.

We found that rapid intermittent hypoxia increases *HIF1A* mRNA in HCT116 cells ((C. Martinez et al., 2019) and Fig 1E) and MCF7 cells when compared to normoxia (Fig 1F). In contrast, when HCT116 or MCF7 cells are exposed to oxygen levels seen in chronically hypoxic tumors, expression of *HIF1A* mRNA decreases when compared to normoxia (Fig 1E and F). We also compared the levels of *HIF1A* mRNA in brain (U251), prostate (PC3) and additional breast (MDA-MB-231) cancer cell lines following exposure to normoxia, chronic and intermittent hypoxia (Supp Fig 1). All cell lines showed a similar decrease in *HIF1A* mRNA in chronic hypoxia and an increase in *HIF1A* mRNA following intermittent hypoxia relative to normoxia. This data indicates that expression of *HIF1A* is controlled differently in intermittent hypoxia when compared to chronic hypoxia.

### Increased *HIF1A* mRNA contributes to HIF-1 activity in intermittent hypoxia

To determine whether the increase in HIF-1α and HIF-target gene expression in intermittent hypoxia is dependent on the increase in *HIF1A* mRNA (Fig 1E and F), HCT116 cells were treated with cycloheximide (CHX), a protein synthesis inhibitor, followed by either chronic or intermittent hypoxia. CHX blocks de novo protein synthesis, therefore HIF-1α levels would primarily reflect protein stabilization and degradation processes. As shown in Fig 1G, CHX reduced HIF-1α protein levels in both chronic and intermittent hypoxia, indicating that HIF-1α accumulation is dependent on continuous protein synthesis in both forms of hypoxia. This also suggests that HIF-1α still undergoes some proteasomal degradation in the presence of either form of hypoxia, though the rate of degradation under individual conditions may vary. CHX also decreased the expression of the downstream HIF-1 target genes, *CA9* (carbonic anhydrase IX) and *HK2* (Supp Fig 2), indicating that de novo HIF-1α protein synthesis is required for HIF-1 activation and HIF-target gene expression to varying degrees in both forms of hypoxia. Based on Fig 1G, the increase in HIF-1α observed in the absence of CHX for both chronic and intermittent hypoxia was due, at least in part, to new protein synthesis.

**Figure 2:**
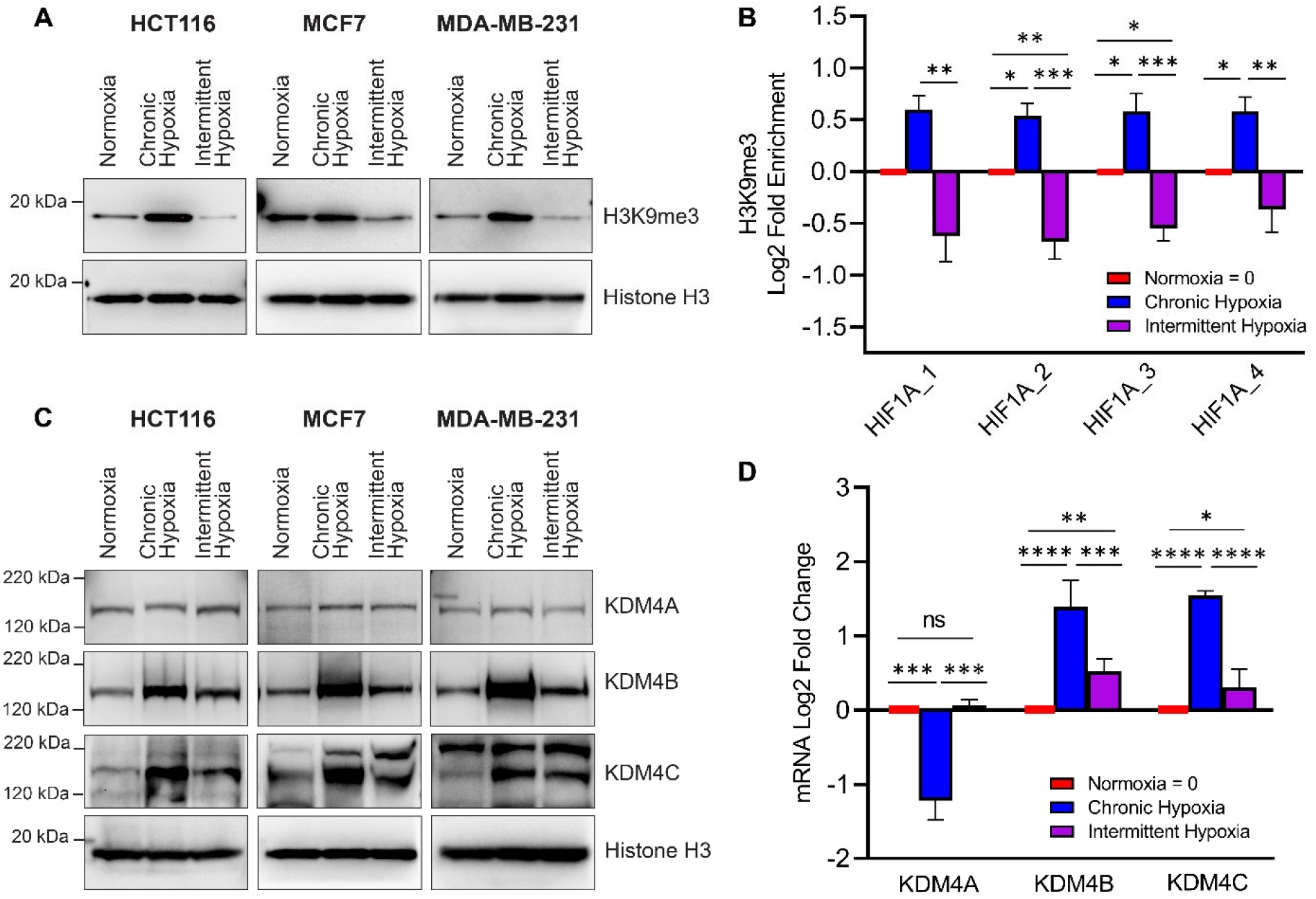
Intermittent hypoxia increases *HIF1A* gene expression by altering histone methylation at the *HIF1A* gene locus. Cells were exposed to normoxia, chronic hypoxia or intermittent hypoxia for 18 h. (A) Western blot of H3K9me3 in HCT116, MCF7 and MDA-MB-231 cells (total protein). (B) ChIP was performed in MCF7 cells for H3K9me3 and enrichment on four different sites along the *HIF1A* gene relative to the total amount of input chromatin was quantified by qPCR. Results are the mean ± S.E.M of 4 independent experiments run in duplicate. Values are normalized to normoxic levels which is set to 0 on the Log2 scale. (C) Western blot of KDM4A, KDM4B, and KDM4C in HCT116, MCF7 and MDA-MB-231 cells (total protein). Total Histone H3 is used as a loading control for (A) and (C). (D) mRNA levels of KDM4A, KDM4B and KDM4C in HCT116 cells measured by qRT-PCR. Values are normalized to normoxic expression which is set to 0 on the Log2 scale. Results are the mean ± S.E.M of 3 independent experiments. ^*^ p < 0.05, ^**^ p < 0.01, ^***^ p < 0.001, ^****^ p < 0.0001. Asterisks above a line compare data points connected by the line.

To further explore HIF-1α protein degradation in intermittent and chronic hypoxia, we treated HCT116 cells with MG262, a proteasome inhibitor. The proteasome is the primary degradation pathway for HIF-1α and by using MG262, we can observe how chronic and intermittent hypoxia affect HIF-1α protein expression without competing degradation processes affecting the final protein quantity. When normoxic HCT116 cells were treated with MG262 (Fig 1H), HIF-1α accumulated due to blocked oxygen-dependent and -independent proteasomal degradation (lane 2). In chronic hypoxia, HIF-1α was stabilized (lane 3) and the addition of MG262 slightly increased HIF-1α protein (lane 4), confirming that low levels of proteasomal degradation still occurs in chronic hypoxia. In intermittent hypoxia, there were lower levels of HIF-1α protein (lane 5) when compared to chronic hypoxia (lane 3), but still more than in normoxia (lane 1, see overexposed blot, Fig 1H). However, when MG262 is added to cells under intermittent hypoxia (lane 6), HIF-1α accumulated well above the observed level in chronic hypoxia, so much so that it was difficult to capture all conditions within the dynamic range of chemiluminescent detection. This indicates that there are higher levels of proteasomal degradation occurring in intermittent hypoxia than in chronic hypoxia, which explains the lower HIF-1α levels in intermittent hypoxia relative to chronic hypoxia. Taken together with the *HIF1A* mRNA (Fig 1E and F) and CHX (Fig 1G) data, this result also suggests that the increase in *HIF1A* mRNA expression in intermittent hypoxia directly contributes to the levels of HIF-1α protein, and when competing degradation processes are inhibited, this leads to a greater increase in HIF-1α (Fig 1H).

These results led us to hypothesize that an increase in *HIF1A* mRNA synthesis in intermittent hypoxia (which does not occur in chronic hypoxia) enables a more robust HIF-1 response to a severe hypoxic event following intermittent hypoxia. To test this hypothesis, we exposed MCF7 cells to 18 hours of normoxia, chronic hypoxia or intermittent hypoxia, that was immediately followed by an additional acute period of chronic hypoxia (4 hours of chronic hypoxia) (total of 22 hours). As expected, 18 hours of normoxia showed no HIF- 1α (Fig 1I, lane 1). The addition of 4 hours of chronic hypoxia following 18 hours of normoxia led to a substantial increase in HIF-1α (lane 2), similar to the amount of HIF-1α present after 18 hours of chronic hypoxia (lane 3). 18 hours of intermittent hypoxia led to a moderate increase in HIF-1α (see overexposed blot lane 4, Fig 1I). However, the largest increase of HIF-1α in any of the conditions occurred when cells were exposed to 18 hours of intermittent hypoxia followed by 4 hours of chronic hypoxia (lane 5). Similar to the MG262 data, this experiment demonstrates that the increased *HIF1A* mRNA in intermittent hypoxia is translated into higher levels of HIF-1α protein, though in intermittent hypoxia some of this is likely degraded via oxygen-dependent degradation. When oxygen-dependent degradation is halted following intermittent hypoxia exposure, via a brief exposure to more severe chronic hypoxia, there is a more robust HIF-1α response due to the increased level of *HIF1A* expression and protein synthesis generated by the preceding intermittent hypoxia.

### Histone 3 lysine 9 methylation increases in chronic hypoxia while decreasing in intermittent hypoxia

We identified that intermittent hypoxia increases *HIF1A* mRNA expression, while chronic hypoxia dampens *HIF1A* mRNA expression, indicating differential regulation depending on the type of hypoxia exposure. The mechanism regulating these differences was unclear. Intermittent hypoxia has been proposed to activate transcription factors NF-κB (Ryan, Taylor, & McNicholas, 2005) and NRF2 (Zhou et al., 2017) and there are binding sites for both NF-κB (Rius et al., 2008) and NRF2 (Lacher, Levings, Freeman, & Slattery, 2018) in the promoter region of the *HIF1A* gene. Therefore, we assessed NF-κB and NRF2 activity in HCT116 cells in intermittent hypoxia using luciferase reporter gene assays. Positive controls (TNF-α and tBHQ) increased NF- κB and NRF2 activity in HCT116 cells, but neither changed in activity following exposure to intermittent hypoxia (Supp Fig 3).

**Figure 3:**
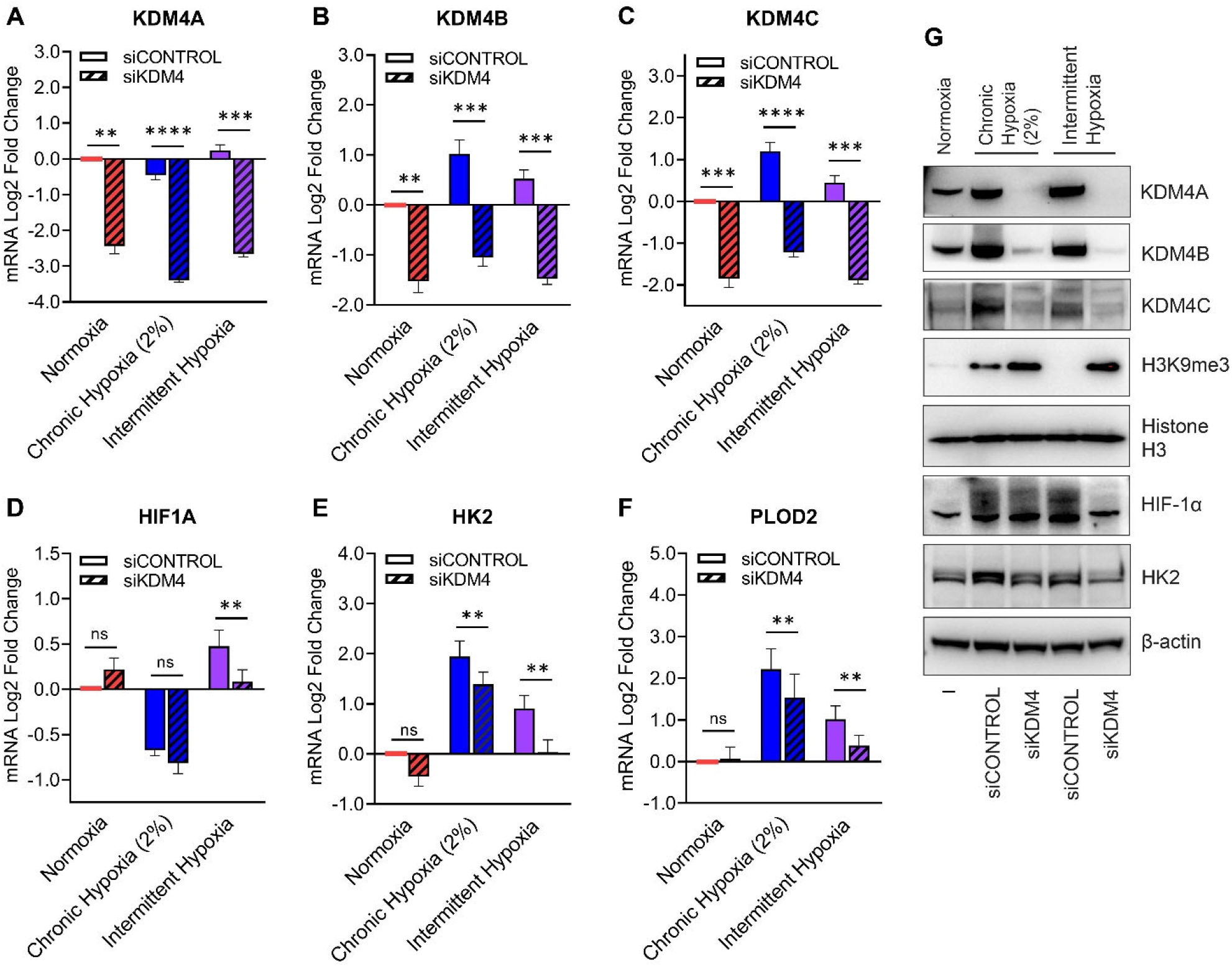
Depletion of KDM4A/B/C increases methylation of H3K9 leading to a decrease in HIF-1α in intermittent hypoxia. MCF7 cells were transfected with combined siKDM4A, siKDM4B, and siKDM4C (referred to as siKDM4 in figure) followed by exposure to normoxia, chronic hypoxia (2%) and intermittent hypoxia for 18 h. mRNA expression of (A) KDM4A, (B) KDM4B, (C) KDM4C, (D) HIF1A, (E) HK2, and (F) PLOD2. All values are normalized to normoxic expression levels which is set to 0 on the Log2 scale. Results are the mean ± S.E.M n = 4 independent experiments. (G) Nuclear extracts of KDM4A, KDM4B, KDM4C, HIF-1α, H3K9me3, and histone H3 (loading control), and cytoplasmic extracts of HK2 and β-actin (loading control).

Negative results for NF-κB and NRF2 led us to explore other potential mechanisms regulating expression of the *HIF1A* gene in intermittent hypoxia. Histone methylation patterns can affect the expression of genes and chronic hypoxia is associated with histone hypermethylation (Hancock et al., 2015), while the effect of intermittent hypoxia on histone methylation has not been investigated. Furthermore, the methylation status of histone 3 lysine 9 (H3K9) has been previously linked to expression of the *HIF1A* gene (Dobrynin et al., 2017), therefore we measured H3K9 trimethylation (H3K9me3) following exposure to normoxia, chronic hypoxia and intermittent hypoxia. In agreement with previous studies (Batie et al., 2019; Dobrynin et al., 2017), we found that H3K9me3 increases in HCT116, MCF7 and MDA-MB-231 cells exposed to chronic hypoxia relative to normoxia (Fig 2A). Surprisingly, and in contrast to chronic hypoxia, we found that intermittent hypoxia decreased levels of H3K9me3 below the levels observed in normoxia (Fig 2A). H3K9me3 is associated with heterochromatin and gene silencing, therefore a global decrease in H3K9me3 induced by intermittent hypoxia is likely resulting in an increase in expression of associated genes (Dobrynin et al., 2017; Hancock et al., 2015). This supports the hypothesis that a reduction in H3K9me3 in intermittent hypoxia may increase expression of *HIF1A*.

### H3K9me3 accumulates at the *HIF1A* gene in chronic hypoxia and decreases in intermittent hypoxia

H3K9me3 has been found to be a repressive marker for *HIF1A* gene expression in chronic hypoxia (Dobrynin et al., 2017). To determine whether the global decrease in H3K9me3 in intermittent hypoxia (Fig 2A) was specifically occurring at the *HIF1A* gene locus, we carried out a chromatin immunoprecipitation qPCR (ChIP- qPCR) assay using MCF7 cells. Following exposure to normoxia, chronic hypoxia and intermittent hypoxia, cells were lysed and H3K9me3 was immunoprecipitated. Co-immunoprecipitated DNA was purified and then subject to qPCR using primers designed to detect *HIF1A* introns. For all regions measured, we found an increased quantity of the *HIF1A* gene in chronic hypoxia compared to normoxia (Fig 2B). This indicates enrichment of H3K9me3 at the *HIF1A* gene in chronic hypoxia and supports previous work done in RKO cells (Dobrynin et al., 2017). In contrast, there was a decrease in immunoprecipitated *HIF1A* in intermittent hypoxia compared to normoxia (Fig 2B), indicating a loss of H3K9me3 at the *HIF1A* gene.

### Histone demethylases KDM4A, -B, and -C increase under intermittent and chronic hypoxia

Histone demethylases (KDMs) alter chromatin by demethylating histones at lysine residues. KDMs belong to the same 2-oxoglutarate-(2-OG)-dependent family of oxygenases as the PHDs and require oxygen to function (Hancock et al., 2015). The KDM4 sub-family removes tri-methyl and di-methyl marks on H3K9 and specific KDM4 enzymes (KDM4B and KDM4C) are transcriptionally upregulated by HIF under hypoxia (Beyer, Kristensen, Jensen, Johansen, & Staller, 2008; Pollard et al., 2008; Xia et al., 2009). Another member (KDM4A) is believed to possess oxygen-sensing capabilities (Hancock et al., 2017), meaning that physiological changes from normoxia to hypoxia inactivate the enzyme. While KDM4A does not appear to be a HIF-target gene, the half-life of KDM4A is prolonged in hypoxia through an unknown mechanism (Black et al., 2015; Dobrynin et al., 2017; Van Rechem et al., 2011), resulting in higher levels of KDM4A in low oxygen conditions, though it is likely to be inactive (Hancock et al., 2017).

We hypothesized that KDM4A may lose activity in chronic hypoxia (increasing H3K9me3), while retaining activity in intermittent hypoxia due to sufficient oxygenation between hypoxic fluctuations, enabling ongoing demethylation of H3K9 similar to normoxia. Furthermore, if intermittent hypoxia increases expression of the HIF-target genes, *KDM4B* and *KDM4C*, as was seen with other HIF-target genes in Fig 1, then this could lead to further H3K9 demethylase activity, even beyond the levels seen in normoxia. To test this hypothesis, we exposed HCT116, MCF7 and MDA-MB-231 cells to normoxia, chronic hypoxia and intermittent hypoxia, and assessed the expression of KDM4A, KDM4B and KDM4C protein and mRNA. We found no changes in the protein levels of KDM4A in cells exposed to normoxia, chronic hypoxia or intermittent hypoxia (Figure 2C). In contrast, protein levels of KDM4B and KDM4C rose substantially in chronic hypoxia and to an intermediate level in intermittent hypoxia (Figure 2C). *KDM4A* mRNA levels decreased in chronic hypoxia, and did not change from normoxia in intermittent hypoxia, supporting data indicating that *KDM4A* is not a HIF-target gene or induced by hypoxia (Hancock et al., 2015) (Fig 2D). Expression of *KDM4B* and *KDM4C* mRNA increased in a similar pattern to the results in Figure 1B-D, where the strongest increase in HIF-target gene expression occurred under chronic hypoxia and a moderate increase in HIF-target gene expression occurred under intermittent hypoxia. This supports the idea that *KDM4B* and *KDM4C* are HIF-target genes (Hancock et al., 2015) (Fig 2D).

### Histone demethylases KDM4A, KDM4B and KDM4C regulate H3K9me3 levels under intermittent hypoxia

Given that oxygen is a required cofactor for KDM4A, KDM4B and KDM4C (or cumulatively KDM4A-C) activity, these enzymes are thought to be largely inactive in chronic hypoxia (Hancock et al., 2015), while intermittent hypoxia may provide sufficient oxygen to maintain activity of KDM4A-C. Furthermore, because the quantities of KDM4B and KDM4C increase in intermittent hypoxia (Fig 2C and D), there may be an overall increase in H3K9 demethylase activity as compared to normoxia. We tested this hypothesis by using a combined siRNA approach to simultaneously knockdown KDM4A, KDM4B, and KDM4C in MCF7 cells. This was followed by measurement of H3K9me3 and *HIF1A* expression. Simultaneous knockdown of KDM4A, KDM4B and KDM4C was highly effective at reducing mRNA and protein levels of each oxygenase (Fig 3A-C, G). While H3K9me3 levels in chronic hypoxia were already high relative to normoxia, knocking down KDM4A, -B, and -C led to a greater increase in the levels of H3K9me3, indicating some residual demethylase activity in chronic hypoxia (Fig 3G). Furthermore, where H3K9me3 was undetectable in intermittent hypoxia, the triple knockdown fully restored H3K9me3, confirming that the methylation status of H3K9 in intermittent hypoxia is dependent on KDM4A-C activity (Fig 3G). As shown in Fig 3D, the triple knockdown had a statistically insignificant effect on *HIF1A* mRNA expression under normoxia and chronic hypoxia. However, in intermittent hypoxia the triple knockdown significantly reduced *HIF1A* mRNA. The triple knockdown also reduced HIF-1α protein levels under intermittent hypoxia, while having no effect on HIF-1α in chronic hypoxia (Fig 3G). We also identified that knockdown of KDM4A-C reduces the mRNA expression of HIF-target genes, *HK2* and *PLOD2* (Fig 3E and F), and HK2 at the protein level (Fig 3G). Whether this occurs through the effects on *HIF1A* expression or through changes in methylation at the *HK2* and *PLOD2* genes themselves is not clear.

### Peripheral intermittent hypoxia combined with spatially distinct core chronic hypoxia enhances overall HIF-1 activity in tumor spheroids

Obstructive sleep apnea (OSA) is associated with higher rates of cancer and cancer mortality (Hunyor & Cook, 2018). The rapid intermittent hypoxia fluctuations associated with OSA have been hypothesized as one of the mechanisms driving cancer progression, and this may be through enhanced activation of HIF-1 in tumor cells, which can drive tumor aggression (Semenza, 2012). In vitro tumor spheroids replicate some aspects of in vivo tumor biology, and they contain both oxygenated, proliferating cells on the exterior and chronically hypoxic cells in the center (Friedrich, Seidel, Ebner, & Kunz- Schughart, 2009). Therefore, we used a spheroid model to explore how the chronic hypoxia present in the avascular tumor center might interact with OSA-driven intermittent hypoxia present in the blood stream, which would be experienced by cells on the tumor periphery. We used this model to observe localized and total HIF activation under varying forms of hypoxia.

HCT116 were transfected with a green fluorescent protein (GFP) reporter linked to the hypoxia response element (HRE) to which HIF-1 binds (Vordermark, Shibata, & Brown, 2001a). HIF-1 mediated expression of GFP was visualized using phase-contrast, fluorescence, and confocal microscopy. As expected, spheroids developed hypoxic centers that activated HIF-1 as indicated by GFP (Fig 4A). As the spheroids grew over 10 days, an increase in hypoxic regions was observed (Fig 4A). GFP expression in the chronically hypoxic spheroid cores was blocked with *HIF1A* siRNA, indicating GFP fluorescence was specific to HIF-1 activity (Fig 4B). The chronic hypoxia generated in the spheroid core increased expression of HIF-target glycolytic proteins, including HK2, LDHA and GLUT1, further validating this model of chronic hypoxia as an activator of HIF-1 (Fig 4C). The degree of glycolytic protein overexpression correlated with the degree of hypoxia observed via GFP (Fig 4A).

**Figure 4:**
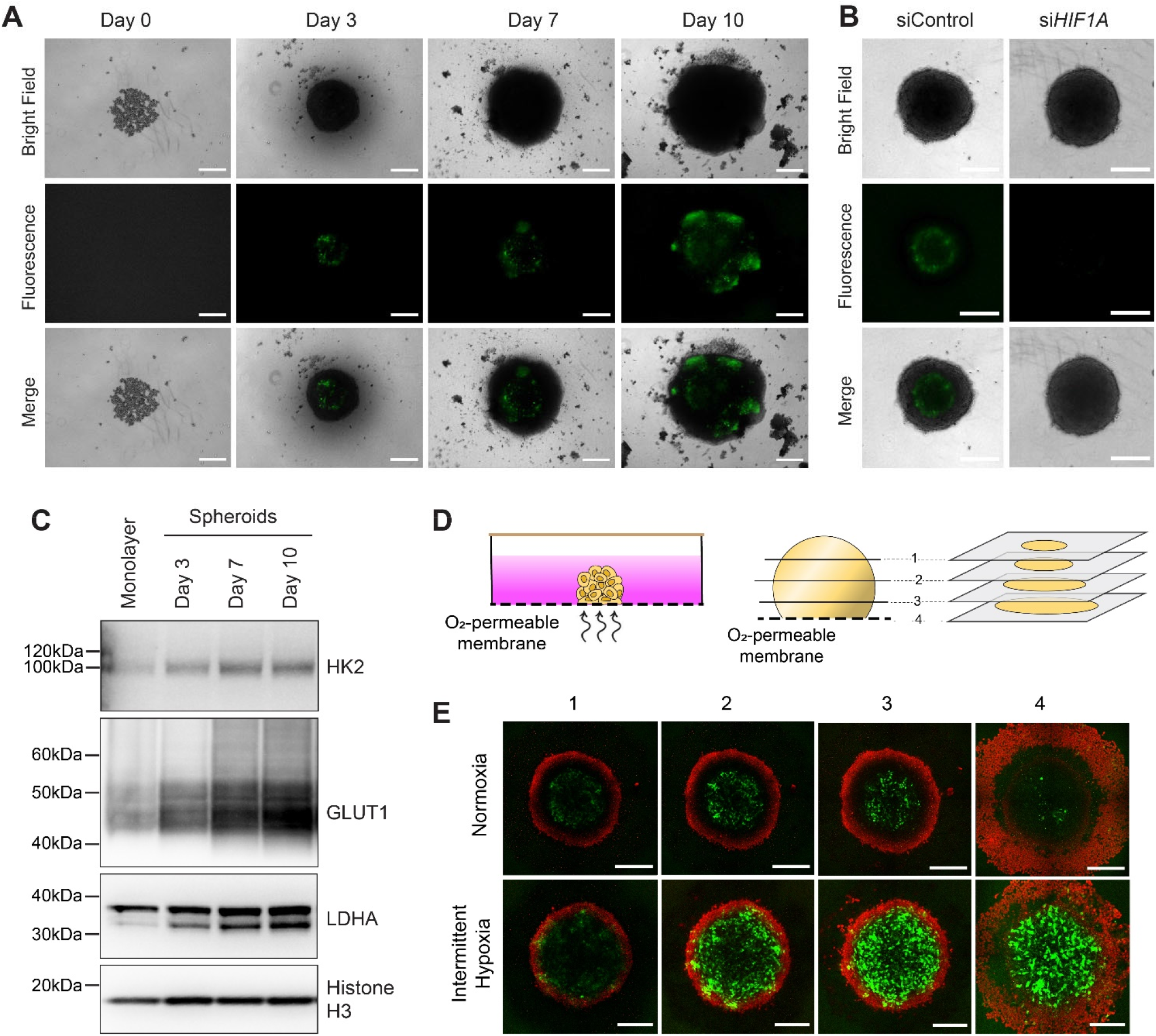
HIF-1 activity in a 3D spheroid model of intermittent hypoxia. HCT116 cells were stably transfected with a 5HRE/EGFP vector and grown as spheroids. (A) Bright field and fluorescence images of HCT116 spheroids expressing GFP after 0, 3, 7, or 10 days of growth. (B) Bright field and fluorescence images of HCT116 spheroids expressing GFP transfected with scrambled control siRNA or *HIF1A* siRNA. (C) Protein expression of HIF-1 target genes, HK2, Glut1, and LDHA in HCT116 cells grown as a monolayer or as spheroids grown over 3, 7, or 10 days. Total histone H3 is used as a loading control. (D) Schematic illustration of spheroid exposure to oxygen conditions using oxygen-permeable membranes; schematic illustration demonstrating how confocal microscopy is used to visualize the spheroid through multiple transverse planes. (E) Confocal fluorescence images of HCT116 cells grown as spheroids over 3 days, transferred onto oxygen-permeable membranes for 24 h, and then exposed to normoxia or intermittent hypoxia over a further 18 h. Red = nuclear red live stain; Green = GFP expression. Scale bar = 300 µm for (A), (B) and (E).

After confirming that in vitro spheroids generate chronic hypoxia near the centre, we wanted to test how intermittent hypoxia stemming from the blood stream might interact with the chronically hypoxic tumour core. 3-day old spheroids were placed onto an oxygen-permeable membrane and left to incubate over 24 h to allow cells closest to the membrane to attach prior to exposure to intermittent hypoxia (Fig 4D). The face of the spheroid at the membrane has direct exposure to supplied oxygen fluctuations which mimics the perfused cells on the tumor exterior. As oxygen must diffuse through the spheroid, this would lead to spatio- temporal gradients of fluctuating oxygen until it reaches the chronically hypoxic regions at the spheroid core. We wanted to explore how HIF-1 activity would be affected by these spatio-temporal changes.

Attached spheroids were exposed to either normoxia or intermittent hypoxia over 18 hours, as with the monolayer model. Immediately after exposure, spheroids were stained with nuclear red live (DNA stain) and confocal images were taken through multiple transverse sections of spheroids in a z-series to examine GFP expression in cells directly attached to the membrane (Figure 4E, Panel 4) compared to cells deeper into the spheroid (Fig 4E, Panel 2 and 3). In normoxic spheroids, there was little to no GFP expression in the cells directly attached to the membrane. Comparatively, GFP expression in the cells directly attached to the membrane, in the spheroids exposed to intermittent hypoxia, was much higher, confirming that HIF-1 is activated by intermittent hypoxia. Surprisingly, while there was increased GFP expression in the hypoxic core of the normoxic spheroids, GFP expression was further increased in the core of the spheroids exposed to intermittent hypoxia. This indicates that cells may respond to a range of spatially and temporally distinct forms of hypoxia through dynamic or overlapping signaling pathways that combine to enhance activity of HIF-1.

## Discussion

Our study reveals a novel oxygen sensing mechanism that is differentially regulated depending on whether cells are exposed to intermittent versus chronic hypoxia (Fig 5). In agreement with previous studies (Hancock et al., 2015), we showed that chronic hypoxia, mimicking oxygen levels in a tumor, inactivates two cellular oxygen sensors, namely the PHDs and KDM4s. This activates the HIF-1 pathway and globally increases H3K9me3 respectively. Traditionally, intermittent hypoxia, including the rapid form which is associated with OSA, is regarded as having the same effect on oxygen sensing pathways as chronic hypoxia. However, our novel results reveal that intermittent hypoxia has different effects on KDM4 activity when compared to chronic hypoxia. KDM4 activity ultimately has consequences for HIF-1 activity, demonstrating a unique relationship between the two oxygen sensing pathways in intermittent hypoxia.

**Figure 5.**
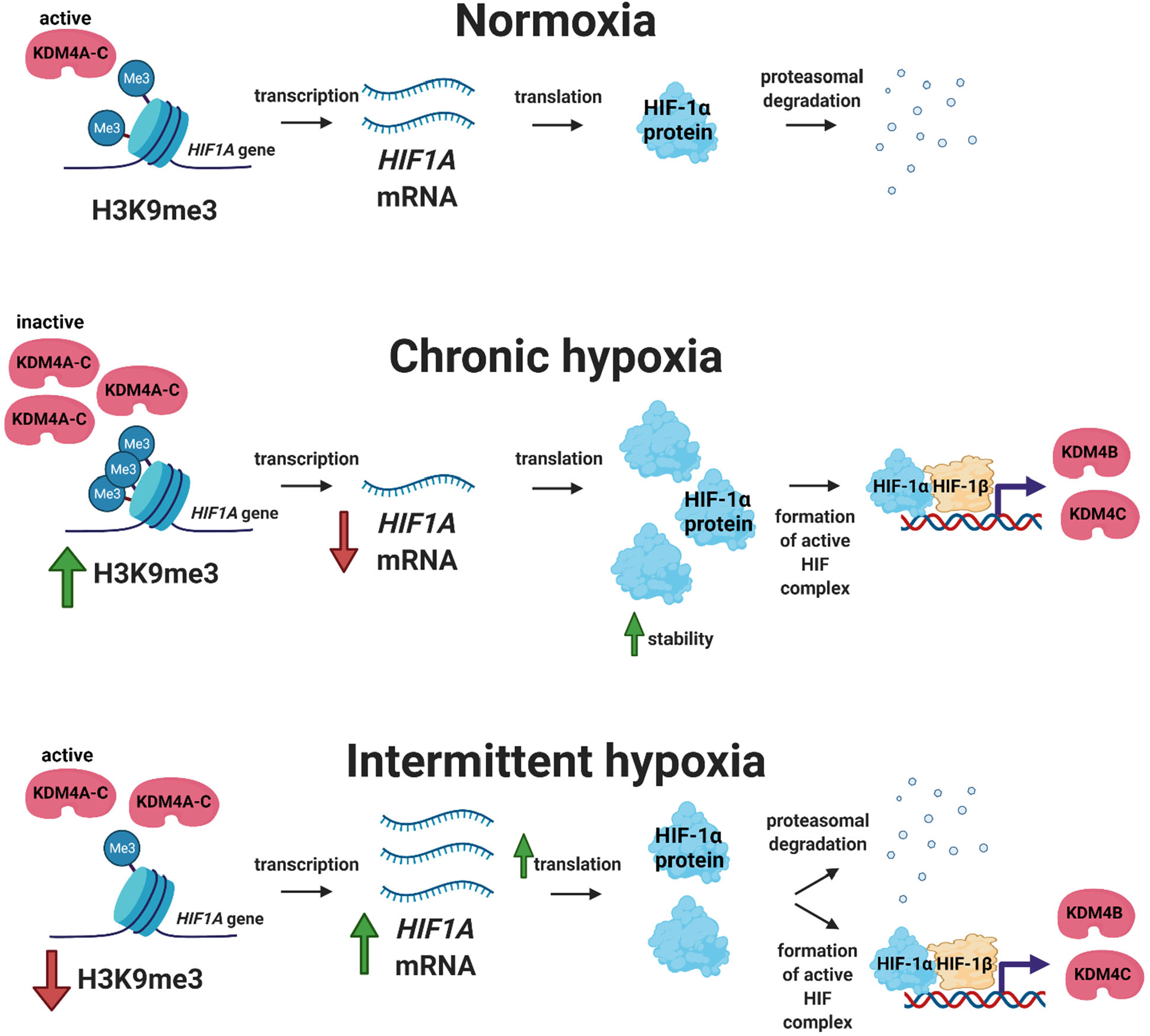
Proposed mechanism of HIF-1 activation in intermittent versus chronic hypoxia. In normoxia, *HIF1A* is constitutively transcribed and translated into HIF-1α, but HIF-1α is post-translationally degraded. In chronic hypoxia, HIF-1α is stabilized, increasing HIF-1 transcriptional activity and the expression of HIF target genes, KDM4B and KDM4C. Despite increased enzyme levels, KDM4A, KDM4B and KDM4C are largely inactive due to limited amounts of oxygen, which are required for KDM activity. This leads to an increase in histone 3 lysine 9 trimethylation (H3K9me3), including at the *HIF1A* locus which ultimately decreases the amount of *HIF1A* mRNA being transcribed. In intermittent hypoxia, HIF-1α increases as compared to normoxia. KDM4B and KDM4C expression levels increase somewhat but not to the same level as chronic hypoxia. However, in contrast to chronic hypoxia, KDM4A, KDM4B and KDM4C activity increases, leading to higher levels of H3K9 demethylation at the *HIF1A* gene than in normoxia or chronic hypoxia. This results in increased *HIF1A* mRNA production.

These results are particularly intriguing when viewed in light of epidemiological, clinical and animal studies that focus on the ‘pre-conditioning effects’ of sleep apnea and cardiovascular disease. Mild to moderate sleep apnea has been suggested to have a protective effect against myocardial infarction in humans and animal models with lower mortality, improved outcomes and reduced infarct size (Gupta et al., 2017; Hu et al., 2020; Mohananey et al., 2017; N. Shah et al., 2013; N. A. Shah, D; Kaplan, R; Yaggi, HK, 2011; Zong et al., 2004). Sleep apnea patients have also been found to have higher levels of serum erythropoietin (EPO) during acute myocardial infarction (Kukwa et al., 2013) and EPO expression is largely controlled by HIF (Haase, 2013). Increased stabilization of HIF is beneficial in myocardial outcomes (Philipp et al., 2006). However, studies indicate that while enhanced expression of HIF-1α is beneficial, protection requires activation *before* the onset of lethal ischemia (Loor & Schumacker, 2008). Our cellular model of intermittent hypoxia mimicking oxygenation changes in sleep apnea induces a significant increase in *HIF1A* mRNA expression, though the full effects of this increase are not immediately reflected at the protein level due to competing HIF-1α degradation controlled by the PHDs. However, when degradation of HIF-1α is halted, either by a drug (MG262, Fig 1G) or possibly through a severe hypoxic insult, such as ischemia or a period of severe hypoxia (Fig 1I), the surplus of *HIF1A* mRNAs are converted into protein, with little degradation, resulting in a corresponding large increase in HIF activity. HIF-1α activation is known to be protective in myocardial infarction, and in this unique situation, an abundance of *HIF1A* mRNA may prove to be beneficial by enabling maximum activation of the HIF-1 pathway. While our studies did not use cardiac or endothelial cells, this could be tested in future using animal models of sleep apnea and myocardial infarction.

Additionally, our spheroid model showed that the HIF-1 response to chronic hypoxia can be further impacted by intermittent hypoxia and may have consequences for tumor biology where tumors are exposed to intermittent hypoxia due to sleep apnea or fluctuations in perfusion. Chronic hypoxia in the spheroid center led to HIF-1 activation, and when combined with intermittent hypoxia on the exterior of the spheroid, HIF-1 activation was greatly enhanced both in the center of the spheroid and closer to the periphery. Several studies have suggested that fluctuating oxygen levels in tumors leads to greater phenotypic variation and a competitive advantage for the tumor (Ardaševa et al., 2020; Gillies, Brown, Anderson, & Gatenby, 2018).

Mathematical modeling of spheroids exposed to fluctuating oxygen predicted that cells on the periphery would have the highest levels of HIF-1 (Leedale et al., 2014), which is somewhat similar to our findings, where the periphery had similar or higher levels of HIF-1 activity when compared to the center. This pathway should be explored further as one potential mechanism linking sleep apnea and tumor aggression and cancer mortality (Hunyor & Cook, 2018).

While HIF pathways have been well studied in acute and chronic forms of hypoxia, with a particular focus on tumors and ischemia, there are much fewer studies on HIF in intermittent forms of hypoxia. New oxygen sensing pathways have been identified very recently, including KDMs (Batie et al., 2019; Chakraborty et al., 2019; Hancock et al., 2017) and ADO (Masson et al., 2019) and there are a very limited number of studies looking at how multiple oxygen sensing pathways might interact under different forms of hypoxia. Our study found that the KDM4 and HIF-1 pathways interact differently depending on whether cells are exposed to intermittent or chronic hypoxia. It is likely that mechanisms and systems have evolved between the oxygen sensing pathways that enable the cell to fine tune the response to hypoxia, whether it is in the form of acute, chronic or intermittent, and based on the severity of hypoxia. One of the limitations to our study thus far is the use of a single pattern of intermittent hypoxia. Studying the biological response to intermittent hypoxia is difficult because the pattern and types of intermittent hypoxia in vivo varies widely and whether or not this response occurs in all forms of intermittent hypoxia is yet to be tested. Future studies will help us to gain further understanding into how new cellular oxygen sensors respond and interact in intermittent hypoxia to generate the hypoxic response.

## Methods

### Cell culture

All cell lines were maintained in media supplemented with 10% fetal bovine serum (FBS), 1x Glutamax, penicillin (50 IU/ml) and streptomycin (50 μg/ml). HCT116 cells were cultured in McCoys 5A media (Gibco). MCF7 and MDA-MB-231 cells were cultured in DMEM media (Gibco). PC3 cells were cultured in DMEM/F12 media (Gibco). Cells were seeded onto oxygen-permeable PDMS membranes 24 h prior to exposure to oxygen conditions. Cell cultures were mycoplasma tested a minimum of four times per year.

### Exposure to oxygen conditions

Exposure to oxygen conditions was carried out as previously described (C.-A. Martinez, Cistulli, & Cook, 2019; C. Martinez et al., 2019). Normoxia was set to 89 mmHg (12% v/v) to reflect tissue oxygenation near major blood vessels (Hunyor & Cook, 2018). Chronic hypoxia was set to 4 mmHg (0.5% v/v) or 15 mmHg (2% v/v) to reflect hypoxia in a tumor. Intermittent hypoxia fluctuated between 5 to 50 mmHg to mimic the *in vivo* oxygen levels previously measured in a tumor in an animal model of OSA (Almendros et al., 2013). Cells were exposed to these oxygen conditions over 18 h unless specified otherwise.

### siRNA

Cells were transfected with siRNA using HiPerfect (Qiagen). We used 20 nM KDM4A FlexiTube GeneSolution siRNA (Qiagen, GS9682), 5 nM siKDM4B (Qiagen, SI00449764), 5 nM siKDM4C (Qiagen, SI05163977), and 20 nM HIF1A Flexitube GeneSolution siRNA (Qiagen, GS3091). Cells incubated for 48 h prior to exposure to oxygen conditions. AllStars Negative control siRNA (Qiagen, SI03650318) was used as a negative control.

### Western blotting

Cells were lysed in RIPA buffer and briefly sonicated. Lysates were mixed in 4x LDS loading buffer containing DTT (ThermoFisher) and heated to 70°C for 10 min. Proteins were separated by SDS-PAGE and resolved using semi-dry transfer. PVDF Membranes were blocked in 5% skim milk. Antibodies used were as follows: HIF-1α (NB100-479, 1:500, Novus), Hexokinase II (TA325030, 1:500, Origene), Glut1 (ab115730, 1:1000, Abcam), LDHA (3582T, 1:1000, Cell Signaling Technologies), JMJD2A (5328, 1:1000, Cell Signaling Technologies), JMJD2B (8639, 1:1000, Cell Signaling Technologies), JMJD2C (ab226480, 1:500, Abcam), Tri- Methyl-Histone H3 (Lys9) (13969, 1:1000, Cell Signaling Technologies), and total Histone H3 (ab1791, 1:2000, Abcam).

### qRT-PCR

Total RNA was extracted using the RNeasy Mini Kit (Qiagen). RNA was quantified using a Nanodrop (Molecular Devices). Total RNA (2 μg) was used as a template for cDNA synthesis using the High-Capacity cDNA Reverse Transcription Kit (Applied Biosystems). For qRT-PCR, pre-designed TaqMan Gene Expression Assays were used with TaqMan Gene Expression Master Mix (LifeTechnologies, ThermoFisher). 20 ng cDNA was used per reaction. qPCR was performed using a QuantStudio 7 Flex Real-Time PCR System. Relative gene expression was normalized to eukaryotic 18S rRNA (Hs03003631_g1) or ACTB (Hs01060665_g1) and fold-change was calculated using the ΔΔCt method. Target gene TaqMan Primers (Assay ID) used for qPCR (ThermoFisher) were as follows: SLC2A1 (Hs00892681_m1), HK2 (Hs00606086_m1), LDHA (Hs01378790_g1), PLOD2 (Hs01118190_m1), P4HA1 (Hs00914594_m1), P4HA2 (Hs00990001_m1), EGLN1 (Hs00254392_m1), EGLN3 (Hs00222966_m1), HIF1A (Hs00153153_m1), KDM4A (Hs00206360_m1), KDM4B (Hs00392119_m1), KDM4C (Hs00909577_m1). Data is expressed as the mean ± S.E.M. from 3 or more independent experiments.

### Spheroids

HCT116 cells were transfected with 5 μg 5HRE/GFP plasmid, a gift from Martin Brown & Thomas Foster (Addgene plasmid #46926; http://n2t.net/addgene:46926 ; RRID:Addgene_46926) (Vordermark, Shibata, & Brown, 2001b) using lipofectamine 3000 (ThermoFisher). Stably expressing cells were selected for using G418 (400 μg/ml), to create a stable HCT116-5HRE/GFP cell line. To form multicellular tumor spheroids, HCT116-5HRE/GFP cells were seeded onto a 96-well clear round bottom ultra-low attachment plate (Costar). On day 3, spheroids were placed onto oxygen-permeable PDMS membranes and incubated for a further 24 h to allow spheroids to attach the membrane prior to exposure to normoxia or intermittent hypoxia.

### Microscopy (confocal and regular fluorescence)

Fluorescence images were acquired using a Zeiss Primovert microscope with 5x and 10x objective. Confocal images were acquired using a Leica TCS SP8 confocal laser scanning microscope with 25xW 0.95 objective. 3D images were collected and stitched using LASX Navigator module. Images were processed and analyzed using Fiji ImageJ (Schindelin et al., 2012).

### Chromatin Immunoprecipitation (ChIP)

Protein was crosslinked to chromatin with 1% formaldehyde for 10 min. Nuclear extracts were prepared by douncing lysates in swelling buffer (25 mM HEPES, pH 8, 1.5 mM MgCl_2_, 10 mM KCl, 0.1% NP-40, 1 mM DTT, EDTA-free protease inhibitors). Nuclei were resuspended in sonication buffer (50 mM HEPES, pH 8, 140 mM NaCl, 1 mM EDTA, pH 8, 1% Triton X-100, 0.5% Na-deoxycholate, 0.1% SDS, EDTA-free protease inhibitors) and sonicated at 70% amplitude for 40 min, (15 s ON, 45 s OFF) at 4°C (QSonica 800R) to shear DNA to an average fragment size of 200 – 1000 bp. Prior to immunoprecipitation, 10% of each lysate was aliquoted to be used as an input reference sample. The remaining lysates were diluted 1:10 in dilution buffer (50 mM Tris-HCl, pH 8, 150 mM NaCl, 2 mM EDTA, pH 8, 1% NP-40, 0.5% Na-deoxycholate, 0.1% SDS, EDTA-free protease inhibitors) and incubated in 10 µg H3K9me3 antibody (1369, Cell Signaling) overnight at 4°C. Immune complexes were captured by incubation with Protein G Dynabeads (Thermo) at 4°C for 2 h. Immunoprecipitates were washed sequentially with a low salt wash buffer (20 mM Tris-HCl, pH 8, 150 mM NaCl, 2 mM EDTA, 1% Triton X-100, 0.1% SDS), a high salt wash buffer (20 mM Tris-HCl, pH 8, 500 mM NaCl, 2 mM EDTA, 1% Triton X-100, 0.1% SDS), a LiCl wash buffer (10 mM Tris-HCl, pH 8, 250 mM LiCl, 1 mM EDTA, 1% NP-40, 1% Na-deoxycholate), followed by TE buffer (10 mM Tris-HCl, pH 8, 1 mM EDTA), and eluted with 120 µL elution buffer (1% SDS, 100 mM NaHCO_3_). Crosslinks were reversed by incubation with 0.2 M NaCl and 10 mg/mL RNase A at 65°C for 4 h. Proteins were digested by incubation with 20 mg/mL proteinase K for 1 h at 60°C. DNA was purified using the Monarch PCR & DNA Cleanup Kit (NEB), according to the manufacturer’s instructions. Immunoprecipitated DNA was quantified by qPCR relative to the total amount of input DNA. Primer sequences for introns on the *HIF1A* gene are described in (Dobrynin et al., 2017).

### Statistics

Statistics were conducted using GraphPad Prism. One-way ANOVA was used when comparing results from normoxia, chronic hypoxia and intermittent hypoxia. Paired two-tailed t-test was used to compare siCONTROL with siKDM4. The number of biological replicates for each experiment are described in the figure legends.

## Acknowledgements

The authors acknowledge the facilities and the scientific and technical assistance of Microscopy Australia at the Australian Centre for Microscopy & Microanalysis at the University of Sydney. Figure 5 was made using BioRender. This study was funded by a Cancer Institute NSW early career fellowship, a University of Sydney fellowship and a NSW Health grant to K.M.C.

## Author contributions

K.M.C. and C.-A.M. conceived the study and analyzed data. C.-A.M and N.B. performed experiments. K.M.C wrote the manuscript with input from C.-A.M. All authors edited the paper and approved the final version.

## Supplemental Information

### Methods: Luciferase reporter gene assays

HCT116 cells were transiently transfected with 5 µg pGL4.32[luc2P/NF-κB-RE/hygro] plasmid (Promega, E8491) or 5 µg pGL4.37[luc2P/ARE/hygro] plasmid (Promega, E3641) using lipofectamine 3000 (ThermoFisher). Cells incubated for 48 h prior to exposure to normoxia or intermittent hypoxia with or without 50 nM TNF-α (positive control for NF-κB activation) or 10 µM tBHQ (positive control for NRF2 activation). Following treatment, cells were lysed in passive lysis buffer (Promega) and mixed with a luciferase assay buffer (100 mM Tris-HCl, pH 7.8, 5 mM MgCl_2_, 0.25 mM Coenzyme A, 0.15 mM ATP) and luciferin-K (final concentration of 471 µM per reaction). Luminescence readings were measured with a reading time of 10 s and a settle time of 10 ms.

**Supplemental Figure 1.**
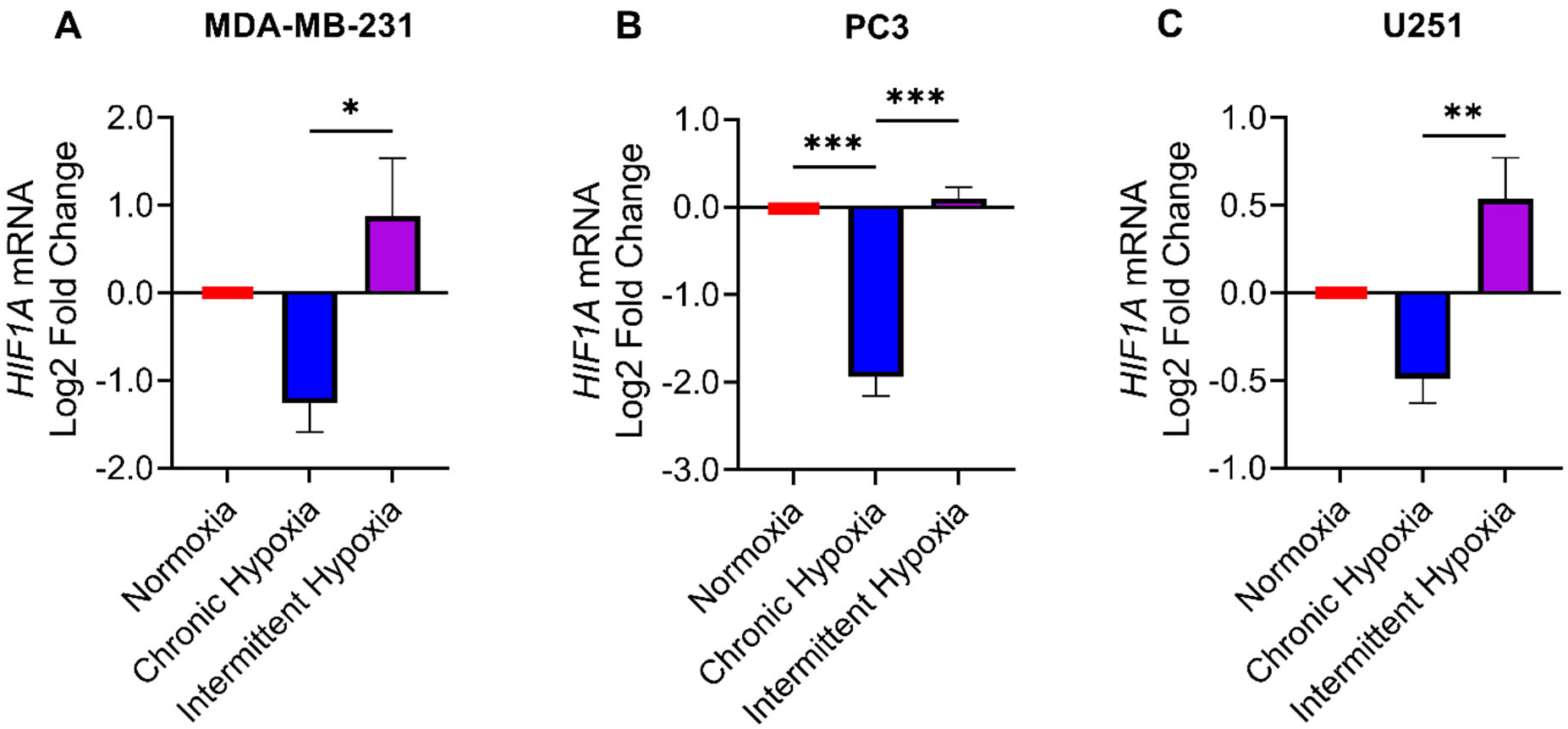
*HIF1A* mRNA expression decreases in chronic hypoxia and increases in intermittent hypoxia in. (A) MDA-MB-231, (B) PC3, and (C) U251 cells after 18 h exposure to normoxia, chronic hypoxia and intermittent hypoxia. All values are normalized to normoxic expression levels which is set to 0 on the Log2 scale (red). Results are the mean ± S.E.M of 3 independent experiments.

**Supplemental Figure 2.**
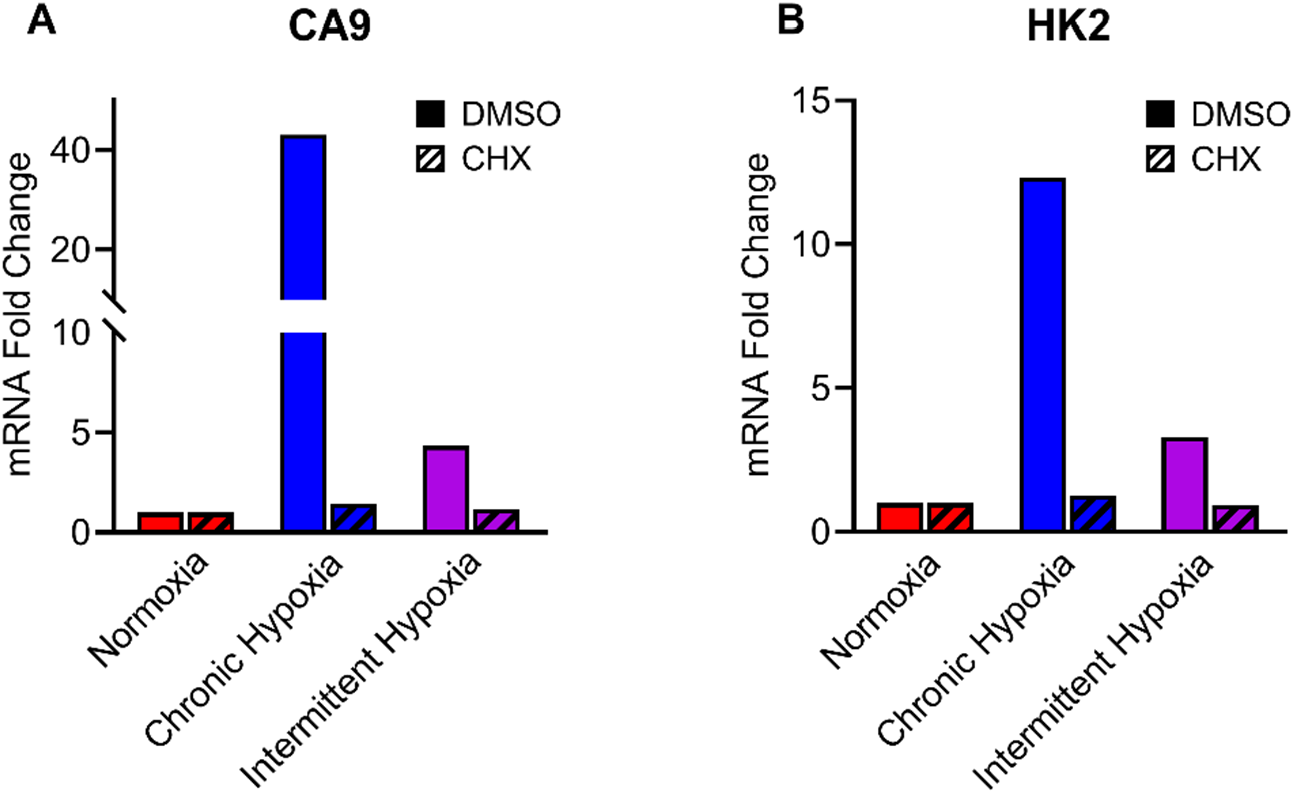
Cycloheximide (CHX) reduces downstream HIF-1 target gene expression in both chronic and intermittent hypoxia in HCT116 cells. (A) CA9 mRNA and (B) HK2 mRNA expression following exposure to normoxia, chronic hypoxia and intermittent hypoxia treated with DMSO or cycloheximide (CHX, protein synthesis inhibitor).

**Supplemental Figure 3.**
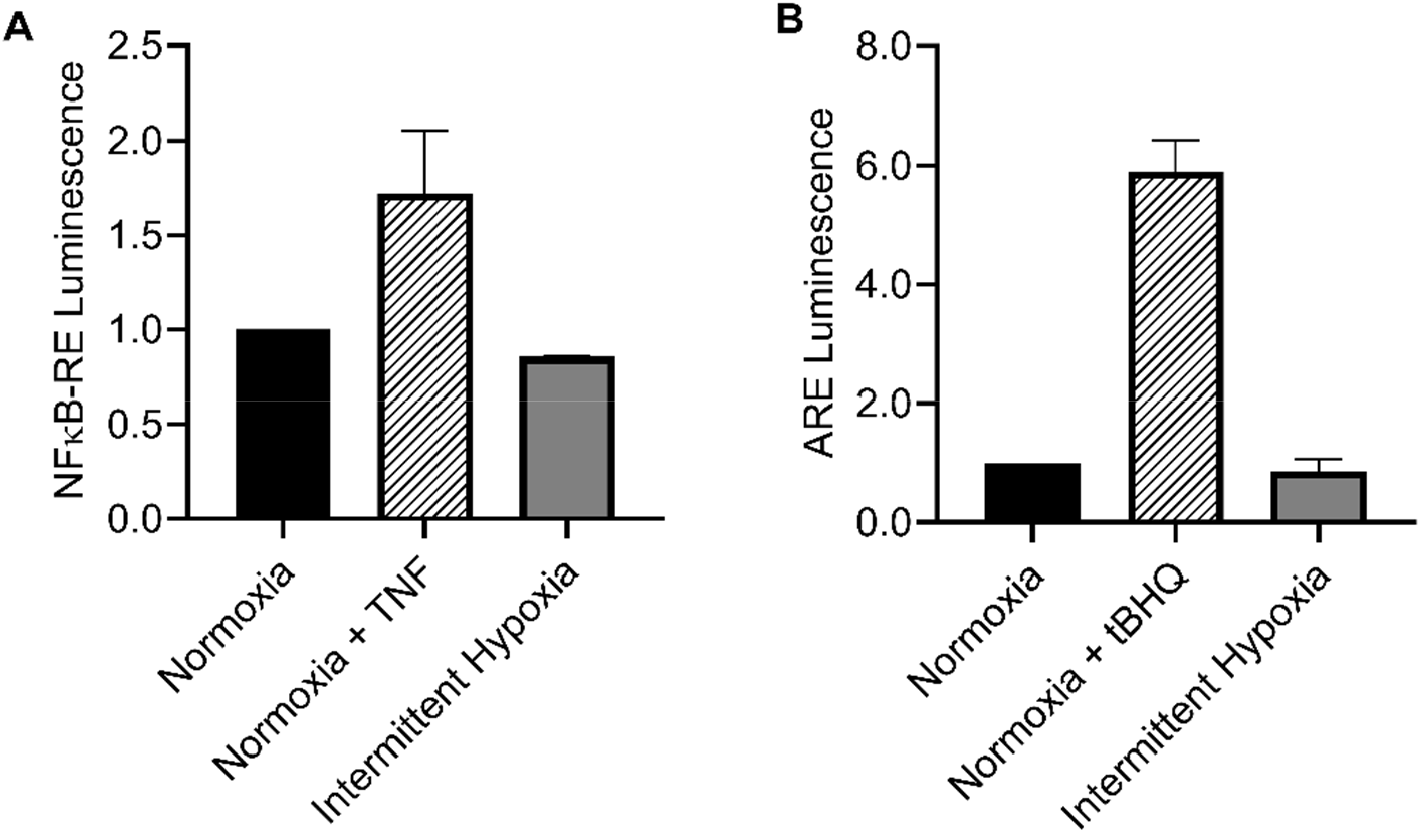
Intermittent Hypoxia does not activate NF-κB or NRF2. (A) NF-κB reporter gene assay in HCT116 cells following 18 h exposure to normoxia, 50 nM TNF-α (tumor necrosis factor-α, NF-κB activator), and intermittent hypoxia. (B) NRF2 reporter gene assay in HCT116 cells following 18 h exposure to normoxia, 10 µM tBHQ (tert-butylhydroquinone, NRF2 activator), or intermittent hypoxia. Results are the mean ± S.E.M of 2 independent experiments.

